# Manipulating Excitation-Inhibition Balance by Temporal Interference Brain Stimulation at Different Frequencies

**DOI:** 10.64898/2026.09.02.748902

**Authors:** Yi-Cheng Fang, Lin Chou, Yao-Yi Tseng, Po-Hsun Chu, Yi-Man Fang, Hai-Yin Chen, Yu-Te Liao, Chih-Hsien Huang, Po-Han Chiang

## Abstract

Temporal interference brain stimulation (TIBS) is a non-invasive neuromodulation approach that can reach deep brain targets by delivering two kilohertz-frequency currents through scalp electrodes, producing a low-frequency amplitude envelope where the fields intersect. Conventional deep brain stimulation suppresses its targets at 130 Hz, and TIBS studies of epilepsy have adopted the same range. However, which neurons TIBS recruits at different frequencies has never been measured. Here, we recorded from genetically defined populations in the mouse hippocampus across envelope frequencies from 10 to 130 Hz, using cell-type-specific fiber photometry, retrograde viral labelling, and immunohistochemistry. A 10 Hz envelope drives both glutamatergic pyramidal neurons and GABAergic interneurons. From 20 Hz onward pyramidal activity falls below baseline while interneuron activity keeps rising, and the two separate maximally at 100 Hz. Retrogradely labelled cortical neurons projecting to CA2 respond weakly and show no frequency dependence, placing the switch inside the local circuit, and c-fos co-staining identifies parvalbumin interneurons as the population recruited at 100 Hz. Overall, this study provides the first in vivo cell-type-resolved evidence for envelope-frequency-dependent neuromodulation and shows that TIBS envelope frequency is a tunable parameter for excitation-inhibition balance. These findings can guide the choice of envelope frequency in the clinical research of TIBS.

## 1 Introduction

Although deep brain stimulation (DBS) with electrode implant is a well-established therapy for several neurological and psychiatric disorders (Lozano et al., 2019; Deuschl et al., 2006; Holtzheimer et al., 2017), fewer than 10 % of patients who meet clinical criteria for DBS ultimately undergo the procedure, largely because of the surgical risks and hardware burden of stereotactic electrode implantation (Lange et al., 2017; Bouthour et al., 2019; Hartmann et al., 2019; Buhmann et al., 2017). Existing non-invasive brain stimulation (NIBS) alternatives cannot fully substitute. Transcranial magnetic stimulation (TMS) reaches <4 cm beneath the scalp before eliciting off-target pain and muscle contraction (Deng et al., 2013; Roth et al., 2007). Transcranial direct- and alternating-current stimulation (tDCS/tACS) sacrifice spatial focality because the current distributes across the whole brain volume, and reaching deep targets suprathresholdly would exceed safety limits (Yavari et al., 2018; Bikson et al., 2016). Transcranial focused ultrasound (FUS) achieves better spatial resolution but is constrained by skull attenuation, beam aberration, and an incompletely characterised mechanism (Legon et al., 2018). Wireless nanoparticle-mediated deep-brain stimulation, in which remotely activated nanotransducers drive deep targets through magnetothermal, magnetomechanical, magnetoelectric, or piezoelectric effects (Hescham et al., 2021; Su et al., 2022; Cheng et al., 2025; Fan et al., 2023), removes the implanted hardware but still requires craniotomy for nanotransducer injection and therefore remains minimally rather than fully non-invasive. A method combining deep penetration, spatial focality, and non-invasiveness within established safety limits would substantially expand the therapeutic and experimental scope of brain stimulation.

Temporal interference brain stimulation (TIBS) addresses this need by delivering two high-frequency (>1 kHz) alternating currents through separate pairs of scalp electrodes at slightly different frequencies, f₁ and f₂ = f₁ + Δf (Grossman et al., 2017). Individual neurons cannot follow the kilohertz carriers due to membrane low-pass filtering, so the carriers themselves do not drive firing; however, at the spatial intersection of the two fields the superposed signal forms an amplitude envelope oscillating at Δf, which can entrain neuronal firing in the targeted deep region while leaving overlying cortex unaffected (Grossman et al., 2017). Envelope frequency appears to determine the direction of the effect. At low Δf, 5 Hz envelope TIBS of the human hippocampus modulates BOLD signals and improves associative memory in healthy volunteers (Violante et al., 2023). At high Δf, 130 Hz envelope stimulation of the mouse hippocampus suppresses pathological fast ripples and interictal epileptiform discharges in a kindling model of epilepsy (Acerbo et al., 2022), and similar suppression of epileptic biomarkers has recently been demonstrated in patients with mesial temporal lobe epilepsy (Missey et al., 2026). Envelope-driving currents ≤2 mA per electrode pair fall within established safety limits for transcranial stimulation (Cassara et al., 2025).

Despite this progress, the cellular mechanism by which TIBS engages neural circuits, and how envelope frequency determines whether stimulation activates or suppresses the target region, remains largely unknown. Four cellular mechanisms have been debated for the suppressive effect of high-frequency conventional DBS: (i) selective recruitment of local GABAergic interneurons that override pyramidal output; (ii) presynaptic synaptic depression or vesicle depletion; (iii) direct depolarisation block of stimulated cell bodies; and (iv) information-lesion-type disruption of firing patterns (Chiken & Nambu, 2016). These mechanisms predict qualitatively different cell-type-level signatures, but the pan-neuronal LFP recordings that have characterised in vivo TIBS to date cannot distinguish them (Acerbo et al., 2022).

A biophysical prediction from computational modelling further motivates a cell-type-resolved approach: envelope demodulation by neurons requires active rectification by voltage-gated ion channels, dominated by fast sodium-channel kinetics, so cell types with different ion-channel compositions should demodulate different envelope frequencies (Mirzakhalili et al., 2020). Pyramidal projection neurons predominantly express Nav1.6 with slow delayed-rectifier potassium channels (Kole et al., 2008), whereas parvalbumin-expressing (PV⁺) GABAergic interneurons combine Nav1.1 with Kv3-family fast-deactivating potassium channels that sustain gamma-frequency firing (Rudy & McBain, 2001; Ogiwara et al., 2007); the two populations are therefore predicted to preferentially demodulate lower and higher envelope frequencies, respectively.

All in vivo TIBS recordings reported so far have used pan-neuronal indicators or local field potentials (Grossman et al., 2017; Violante et al., 2023; Acerbo et al., 2022). These signals sum activity across cell types, so a fall in pyramidal firing combined with a rise in interneuron firing cannot be distinguished from a uniform reduction in both. A prior study demonstrates cell-type-resolved data in acute cortical slices at a 20 Hz envelope (Caldas-Martinez et al., 2024). However, conventional DBS of hippocampal-circuit targets runs at 130 to 145 Hz (Fisher et al., 2010; Lozano et al., 2016), and the TIBS protocol that suppressed epileptic biomarkers in the hippocampus used a 130 Hz envelope (Acerbo et al., 2022). No cell-type-specific measurement has been made in that range in any preparation.

Here, we combine cell-type-specific fiber photometry with Thy1-GCaMP6s (Dana et al., 2014) spatial validation and c-fos / parvalbumin double immunostaining to map TIBS-evoked neuronal activity in the mouse hippocampus, a target with direct translational relevance to the recent human TIBS trial (Violante et al., 2023), across envelope frequencies from 10 to 130 Hz. We find that 10 Hz envelope stimulation activates both glutamatergic and GABAergic populations, but from 20 Hz onward TIBS actively suppresses glutamatergic pyramidal activity below baseline while progressively recruiting GABAergic interneurons, with the divergence between the two populations reaching a maximum at 100 Hz. Retrogradely labelled entorhinal and ectorhinal cortex neurons projecting to CA2 show a milder, monotonically declining response, suggesting that the frequency-dependent E/I crossover is most pronounced in local hippocampal cell types. This pattern is confirmed by c-fos and parvalbumin co-staining. Our results redefine TIBS as a frequency-tunable modulator of hippocampal excitation–inhibition balance, with envelope frequencies above approximately 20 Hz progressively biasing the balance towards inhibition. These findings provide a mechanistic framework for the rational design of TIBS protocols in disorders characterised by E/I dysregulation.

## 2 Results

### 2.1 Validation of TIBS system in dorsal hippocampus

To deliver temporally interfering currents to the deep brain without engaging overlying cortex, we built a custom multi-channel current stimulator that generated two pairs of high-frequency (2.00 kHz) sinusoidal currents whose envelope oscillated at the difference frequency Δf (Figure 1A; Chou et al., 2025). The stimulator’s LabVIEW interface allowed programmable control over the carrier frequencies, envelope frequency, current amplitude, and the duration of the onset and offset amplitude ramps (Figure 1B); in all experiments below, TIBS was delivered with a 2 s linear ramp-up to suppress stimulation onset artefacts (Grossman et al., 2017). The four stimulating electrodes were implanted on the right hemisphere at bregma coordinates spanning anterior to posterior cortex, with a single ground electrode placed contralaterally.

**Figure 1.**
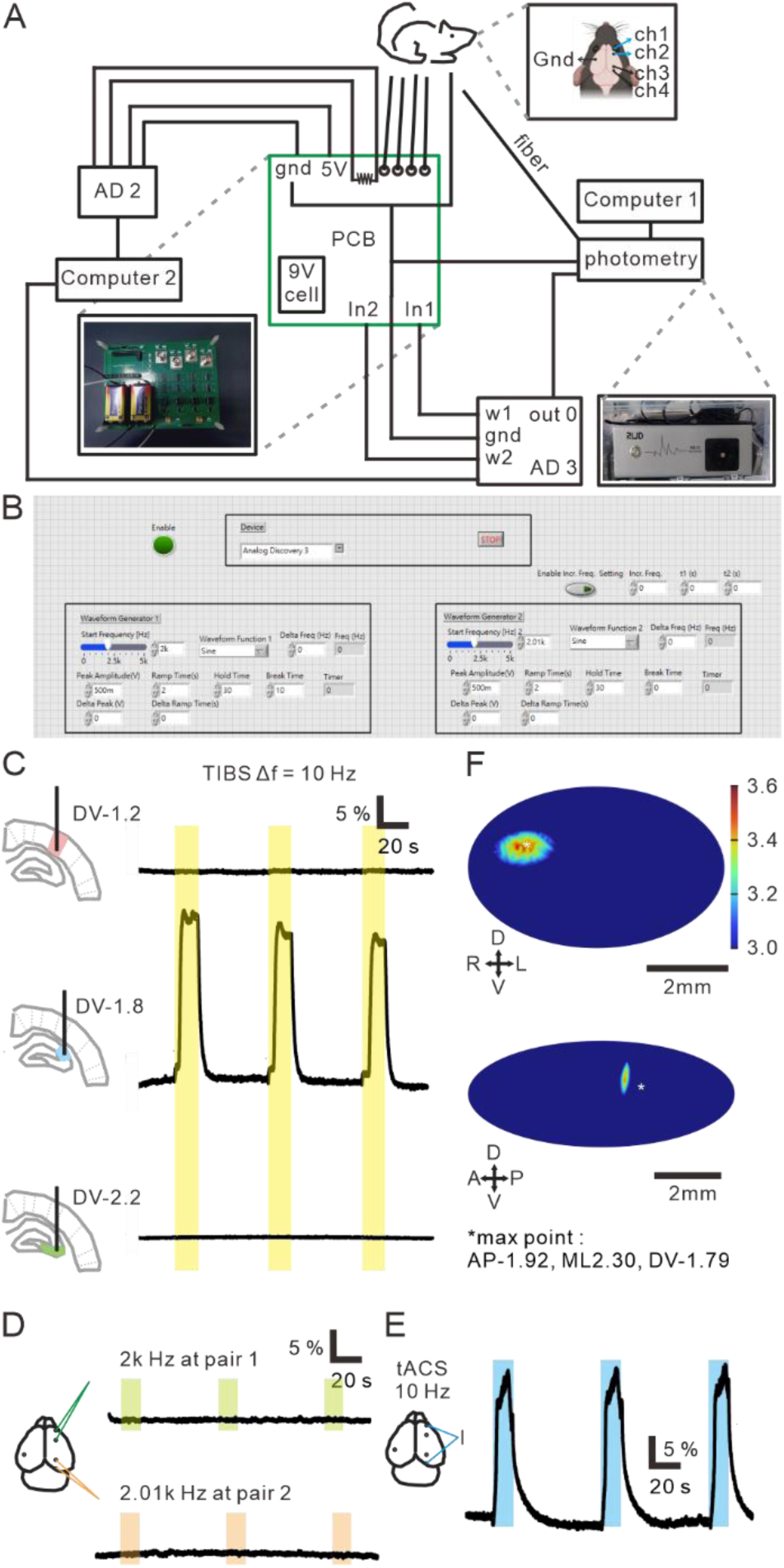
TIBS system, COMSOL simulation, and spatial specificity in dorsal hippocampus. (**A**) Schematic of the custom TIBS stimulation system and experimental setup. Stimulation waveforms were generated in LabVIEW, transmitted through AD3, and converted into biphasic constant-current outputs using a custom PCB-based V-to-I converter. Stimulation currents were delivered through two electrode pairs implanted on the mouse skull. AD2 continuously monitored the stimulation waveforms in real time through current-sensing resistors on the PCB. AD3 additionally generated a digital trigger signal that synchronised fiber photometry acquisition with stimulation onset. The lower panel shows the complete experimental setup, including the custom PCB, stimulation hardware, fiber photometry system, and electrode configuration on the mouse head. (**B**) LabVIEW graphical user interface used for stimulation-parameter control and waveform generation. (**C**) Calcium responses evoked by TIBS in Thy1-GCaMP6s mice at different recording depths, measured by fiber photometry, from the cortex (DV = −1.2 mm), hippocampal CA2 (DV = −1.8 mm), and hippocampal CA3 (DV = −2.2 mm) at AP = −1.94 mm and ML = 2.36 mm. Yellow shaded regions indicate stimulation periods; insets illustrate the recording locations. (**D**) COMSOL simulation of the maximum electric field induced by temporal interference stimulation in the mouse brain, shown in the horizontal (xy), sagittal (yz), and coronal (xz) planes, demonstrating focal modulation around the hippocampal CA2 region. (**E**) Negative-control experiments using high-frequency stimulation delivered through a single electrode pair (top, Pair 1 at 2.00 kHz; bottom, Pair 2 at 2.01 kHz), with calcium activity recorded from hippocampal CA2; insets illustrate the active electrode pair. (**F**) Positive-control experiment using tACS (10 Hz), with calcium activity recorded from hippocampal CA2 while stimulation was delivered between two distal electrodes from different pairs.

To validate the spatial specificity in vivo, we performed fiber photometry at different depth along the implantation track in Thy1-GCaMP6s transgenic mice, which express GCaMP6s under the Thy1 promoter in glutamatergic neurons across cortex and hippocampus (Dana et al., 2014). The optic fiber was advanced through successive cortical and subcortical depths during a single anaesthesia session, and ΔF/F responses to 10 Hz envelope TIBS were measured at each depth (Figure 1C). Calcium responses in the overlying neocortex (DV ∼1.2 mm) were minimal, increased markedly upon entering the dorsal hippocampus at the level of CA2 (DV ∼1.8 mm), and disappeared at deeper levels within CA3 (DV ∼2.2 mm). The peak depth (DV −2)of the in vivo response matched the location of maximum envelope modulation depth predicted by COMSOL Multiphysics (ML2.30 AP-1.92 DV-1.79) (Figure 1D), confirming focal CA2 targeting.

To verify that the CA2 calcium response was generated by the field-intersection envelope rather than by either kilohertz carrier on its own, we recorded three additional control conditions with the fiber positioned at the CA2 depth. Activating only the first electrode pair (carrier f₁ = 2.00 kHz alone) or only the second pair (carrier f₂ = 2.01 kHz alone) evoked no detectable ΔF/F response in CA2 (Figure 1E). In contrast, when transcranial alternating current stimulation (tACS) at 10 Hz was delivered at the same current intensity, a clear calcium response was observed (Figure 1F). These controls confirm that the response evoked by TIBS arises from the envelope formed at the spatial intersection of the two kilohertz fields, aligned with prior TIBS study (Grossman et al., 2017).

### 2.2 Pan-neuronal hippocampal activity attenuates monotonically with TIBS envelope frequency

Having established the spatial specificity of TIBS in dorsal CA2, we next asked how envelope frequency shapes hippocampal activity at the pan-neuronal level during stimulation. To obtain a soma-targeted readout of all neurons in dorsal CA2 that is largely free of neuropil and axonal contributions, we expressed a soma-restricted version of jGCaMP8m under the pan-neuronal synapsin promoter (AAV-hSyn-soma-jGCaMP8m) and recorded photometry during TIBS at six envelope frequencies (10, 20, 40, 70, 100, and 130 Hz), with three stimulation epochs per frequency averaged for each animal (Figure 2A-B). The maximum stimulus-evoked ΔF/F response declined monotonically with increasing envelope frequency, from 3.77 ± 1.70 at 10 Hz to 2.14 ± 0.75 at 130 Hz (Figure 2C, D). Activity changed most steeply between 10 and 70 Hz and plateaued at higher envelope frequencies. An equivalent monotonic decline was observed in a separate cohort expressing the non-soma-restricted hSyn-jGCaMP8m indicator. However, pan-neuronal indicators cannot distinguish uniform attenuation from opposing cell-type-specific changes that partially cancel-out each other, we next recorded from glutamatergic and GABAergic populations separately under the same TIBS protocol.

**Figure 2.**
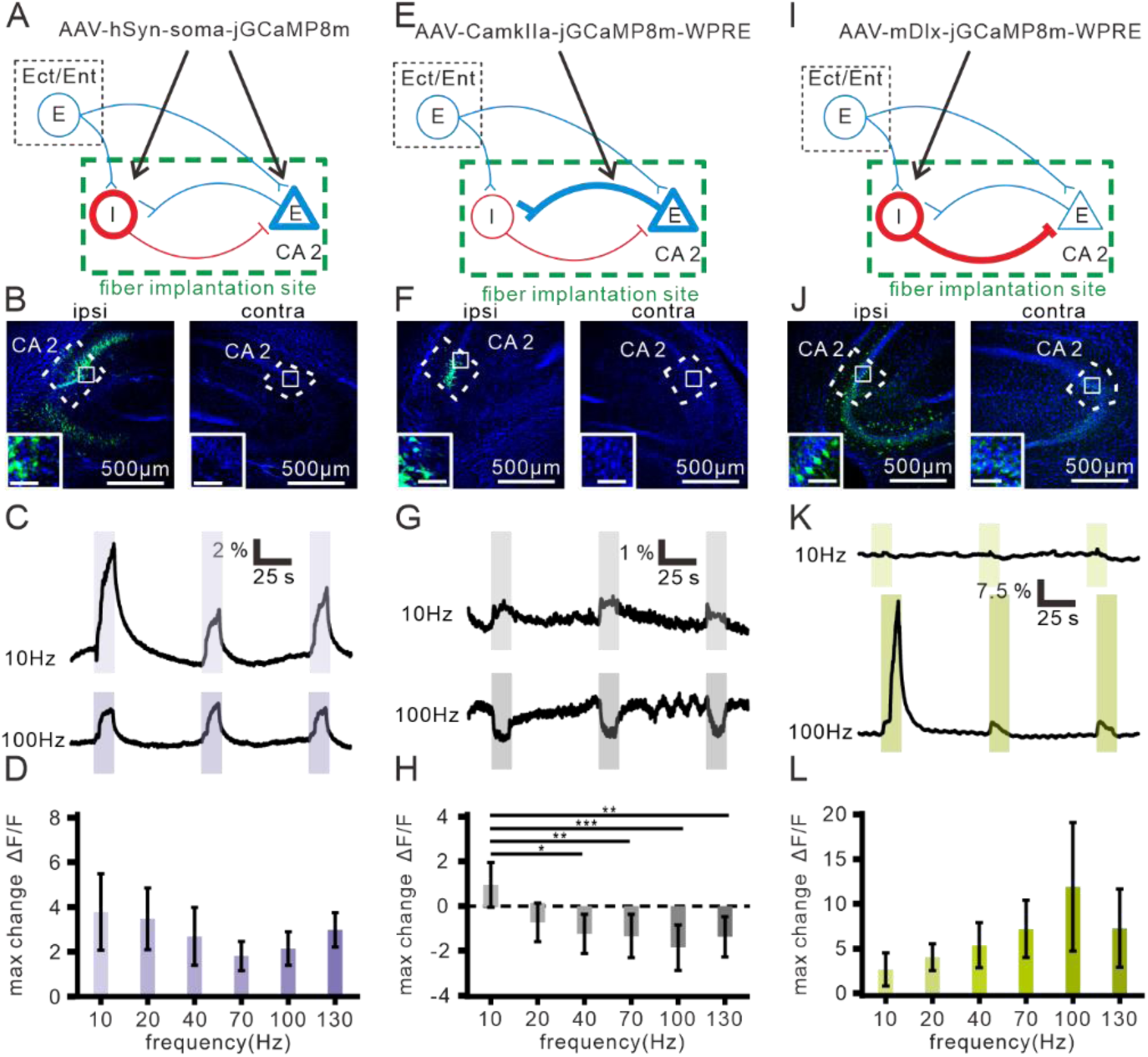
Cell-type-specific calcium responses to TIBS across envelope frequencies in dorsal CA2. (**A**) Schematic of viral expression and fiber photometry recording for soma-targeted pan-neuronal recording (AAV-hSyn-soma-jGCaMP8m). Red and blue cells denote inhibitory and excitatory neurons, respectively; thick lines denote sites of viral expression; dashed boxes denote brain regions; thick green boxes denote the recording location. (**B**) Representative fluorescence images of viral expression in the contralateral (left) and ipsilateral (right) hippocampal CA2. Insets show magnified views of the white-boxed regions. Green, GCaMP8m; blue, DAPI. Scale bars: main, 500 μm; insets, 80 μm. (**C**) Representative calcium traces recorded during 10 Hz (upper trace; 2.00 and 2.01 kHz carriers) and 100 Hz (lower trace; 2.00 and 2.10 kHz carriers) envelope TIBS. Purple shaded regions indicate the stimulation periods. (**D**) Quantification of the calcium response at different envelope frequencies (n = 6). (**E**) Schematic of viral expression and fiber photometry recording for excitatory pyramidal recording (AAV-CaMKIIα-jGCaMP8m-WPRE); similar to (A). (**F**) Similar to (B), but with AAV-CaMKIIα-jGCaMP8m-WPRE injection for expressing GCaMP on pyramidal neurons. (**G**) Similar to (C), but recorded from CaMKIIα-positive pyramidal neurons; gray shaded regions indicate the stimulation periods. (**H**) Quantification of the calcium response (n = 6). (**I**) Schematic of viral expression and fiber photometry recording for inhibitory (GABAergic) recording (AAV-mDlx-jGCaMP8m-WPRE); similar to (A). (**J**) Similar to (B), but with AAV-mDlx-jGCaMP8m-WPRE injection for expressing GCaMP on GABAergic interneurons. (**K**) Similar to (C), but recorded from mDlx-positive interneurons; green shaded regions indicate the stimulation periods. (**L**) Quantification of the calcium response (n = 9). The maximum change of ΔF/F corresponding to the larger magnitude of either the maximum positive or minimum negative deflection during stimulation. TIBS was delivered at 1.9 mA. *p < 0.05, **p < 0.01, ***p < 0.001, Repeated-measures ANOVA followed by Holm’s post-hoc test.

### 2.3 Cell-type-specific photometry reveals a frequency-dependent E/I crossover in dorsal CA2

We first expressed jGCaMP8m selectively in CaMKIIα-positive pyramidal cells of dorsal CA2 (AAV- CamKIIa-jGCaMP8m-WPRE) and recorded photometry under the same TIBS protocol (Figure 2E-F). At 10 Hz envelope frequency, TIBS evoked a positive ΔF/F response (+0.95 ± 1.00). From 20 Hz onward, however, the ΔF/F response inverted and became negative, indicating that the maximum signal during stimulation fell below the pre-stimulation baseline (Figure 2G-H). This suppression progressed monotonically with envelope frequency from −0.73 ± 0.87 at 20 Hz to −1.24 ± 0.87 at 40 Hz and −1.34 ± 0.97 at 70 Hz, reaching its strongest magnitude of −1.87 ± 1.03 at 100 Hz before partially recovering at 130 Hz (−1.37 ± 0.90; Figure 2H). Thus, CaMKII-positive pyramidal activity inverts between 10 and 20 Hz envelope frequency: low envelope frequencies activate this population, whereas envelope frequencies of 20 Hz and above suppress it below baseline, with maximum suppression at 100 Hz.

To examine the contribution of GABAergic interneurons, we expressed jGCaMP8m under the mDlx promoter (AAV-mDlx-jGCaMP8m-WPRE), which selectively drives expression in cortical and hippocampal GABAergic interneurons (Dimidschstein et al., 2016), and recorded photometry under the same six-frequency TIBS protocol (Figure 2I-J). In contrast to the inversion observed in pyramidal neurons, GABAergic interneurons exhibited a positive, monotonically increasing ΔF/F response across the envelope frequency range. The maximum ΔF/F rose from 2.63 ± 1.85 at 10 Hz to 4.04 ± 1.50 at 20 Hz, 5.37 ± 2.55 at 40 Hz, and 7.20 ± 3.20 at 70 Hz, reaching a peak of 11.91 ± 7.20 at 100 Hz before partially declining to 7.28 ± 4.38 at 130 Hz (Figure 2K-L). Thus, while pyramidal neurons were suppressed at envelope frequencies ≥ 20 Hz, GABAergic interneurons were progressively activated across the same range, with both populations reaching their respective extrema at 100 Hz. Comparing the CaMKII and mDlx responses across envelope frequencies (Figure 2L, 2H) shows that the two populations diverge, with a crossover near 20 Hz where the excitatory response falls below baseline, and a maximum divergence of approximately 14 % of ΔF/F at 100 Hz. These results indicate that TIBS envelope frequency shifts hippocampal excitation/inhibition balance, with a crossover near 20 Hz and maximum inhibitory bias at 100 Hz.

### 2.4 High-frequency TIBS preferentially recruits parvalbumin-positive interneurons

To independently verify the cell-type-specific activation patterns observed by photometry, we performed c-fos immunostaining with parvalbumin (PV) co-labelling in wild-type mice that received TIBS at either 10 Hz or 100 Hz envelope frequency and were sacrificed 90 min after stimulation to capture peak c-fos expression (Figure 3). c-fos⁺ cells were counted within predefined regions of interest of the dorsal hippocampus (CA1, CA2, CA3, dentate gyrus) and the overlying primary somatosensory (S1BF) and retrosplenial (RSG) cortex, normalised to DAPI⁺ cell density in the same region, and compared between the ipsilateral (stimulated) and contralateral hemispheres (Figure 3C, H). At 10 Hz envelope stimulation, c-fos⁺/DAPI⁺ density in the ipsilateral CA2 was significantly elevated relative to every other examined region (p = 0.007; One-way ANOVA), including the adjacent CA1, CA3, dentate gyrus, and the overlying cortical fields (Figure 3A-C), consistent with the previously reported hippocampal-selective activation of TIBS (Grossman et al., 2017; Acerbo et al., 2022). At 100 Hz envelope stimulation, ipsilateral CA2 c-fos density showed a similar trend towards elevation but the regional difference did not reach statistical significance (p = 0.327; One-way ANOVA) (Figure 3F-H). Co-staining for PV, a marker that labels fast-spiking GABAergic interneurons predicted to preferentially demodulate higher envelope frequencies (Mirzakhalili et al., 2020), revealed the cell-type origin of these regional patterns. As expected for a constitutive cell-type marker, the density of PV⁺ cells was similar across regions and between the 10 Hz and 100 Hz conditions. Two complementary indices of PV recruitment were then quantified. The fraction of PV interneurons that co-expressed c-fos (c-fos⁺PV⁺ / total PV⁺), which indexes the proportion of the PV population that was active during stimulation, showed no regional difference at 10 Hz envelope frequency (p = 0.376; One-way ANOVA) but became significantly elevated in ipsilateral CA2 relative to all other examined regions at 100 Hz (p = 0.003; One-way ANOVA) (Figure 3D, I). The reciprocal ratio, the fraction of c-fos⁺ cells that co-expressed PV (c-fos⁺PV⁺ / total c-fos⁺), similarly showed no regional difference at 10 Hz (p = 0.449; One-way ANOVA) and a trend towards elevation in ipsilateral CA2 at 100 Hz that did not reach statistical significance (p = 0.355; One-way ANOVA) (Figure 3E, J). Together, these data indicate that 10 Hz TIBS drives predominantly pyramidal activation in CA2, whereas 100 Hz TIBS shifts CA2 activation toward selective PV interneuron recruitment.

**Figure 3.**
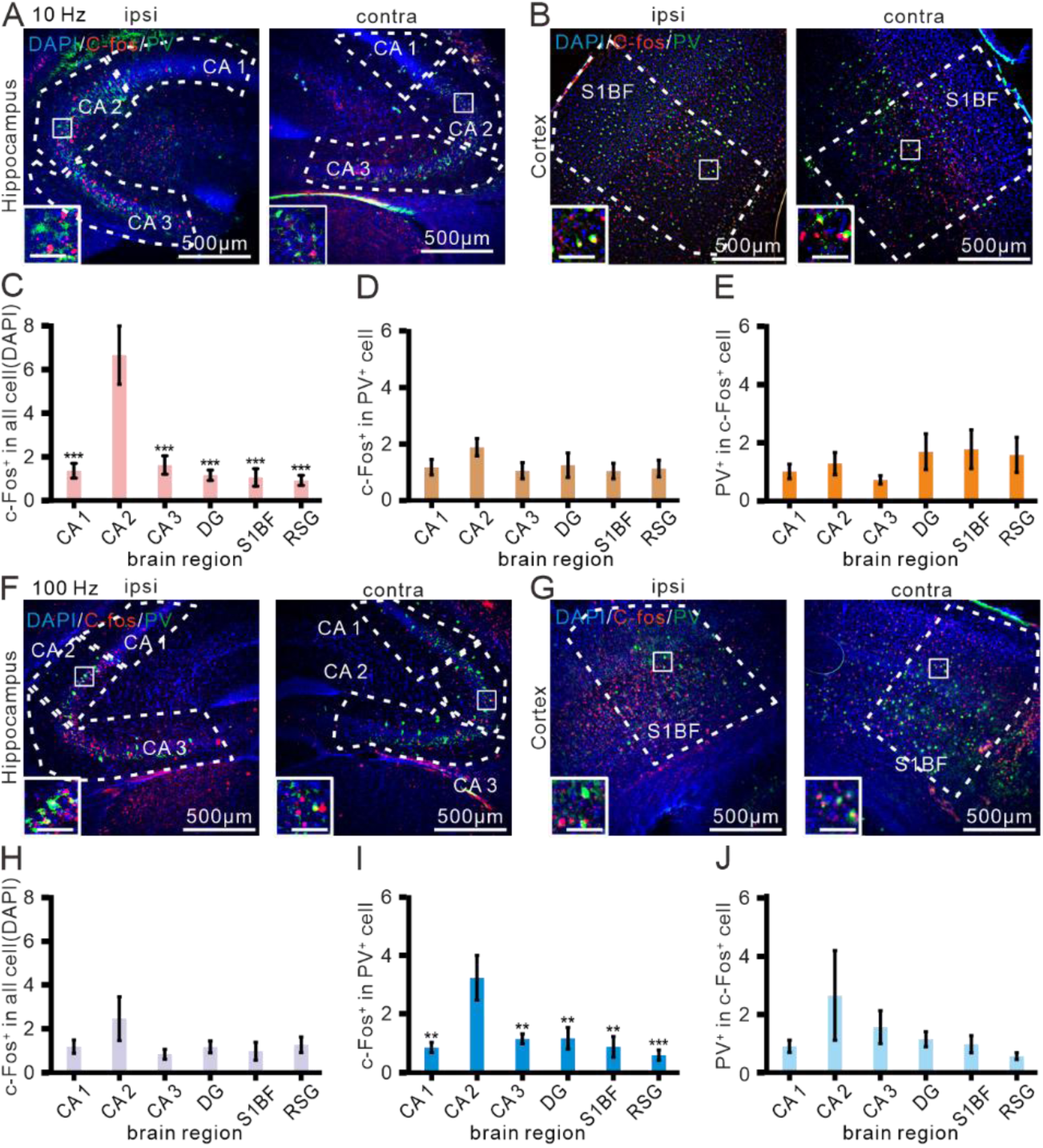
c-fos and PV immunostaining following 10 Hz and 100 Hz TIBS in dorsal CA2. (**A**) Representative immunofluorescence images of c-Fos and PV immunostaining in the ipsilateral (ipsi) and contralateral (contra) hippocampus (CA1, CA2, and CA3) following 10 Hz TIBS. Insets show magnified views of the white-boxed area. Blue, DAPI; green, PV; red, c-Fos. Scale bars: main, 500 μm; insets, 80 μm. (**B**) Representative immunofluorescence images of the ipsilateral and contralateral S1BF following 10 Hz TIBS; insets and scale bars as in (A). (**C**) Quantification of the percentage of c-Fos-positive/DAPI-overlapping area relative to the total DAPI-stained area across CA1, CA2, CA3, DG, S1BF, and RSG following 10 Hz TIBS (pink). (**D**) Quantification of the proportion of c-Fos/PV double-positive cells relative to the total PV cell count across the indicated regions following 10 Hz TIBS (brown). (**E**) Quantification of the proportion of c-Fos/PV double-positive cells relative to the total c-Fos-positive cell count across the indicated regions following 10 Hz TIBS (orange). (**F**) Similar to (A), but following 100 Hz TIBS. (**G**) Similar to (B), but following 100 Hz TIBS. (**H**) Similar to (C), but following 100 Hz TIBS (purple). (**I**) Similar to (D), but following 100 Hz TIBS (azure). (**J**) Similar to (E), but following 100 Hz TIBS (purple). Unless otherwise indicated, TIBS was delivered at 1.9 mA. *p < 0.05, **p < 0.01, ***p < 0.001, Repeated-measures ANOVA followed by Holm’s post-hoc test.

### 2.5 TIBS envelope frequency entrains hippocampal calcium oscillations

To assess whether TIBS modulates the temporal structure of hippocampal activity in addition to its mean amplitude, we recorded photometry signals at a higher sampling rate (150 frames per second) capable of resolving oscillations at the envelope frequencies tested, and computed time-frequency spectrograms and power spectral density (PSD) of the ΔF/F traces during baseline and stimulation for the soma-restricted pan-neuronal (hSyn-soma), CaMKII-positive pyramidal, and mDlx-positive GABAergic interneuron populations (Figure 4). Because the indicator kinetics of jGCaMP8m limit the temporal resolution of calcium-based oscillatory measurements, envelope-frequency entrainment could only be reliably resolved at the lower envelope frequencies (10 Hz and 20 Hz) and was difficult to detect at 40 Hz; higher envelope frequencies (≥70 Hz) lie beyond the effective bandwidth of the indicator and were therefore excluded from the spectral analysis.

**Figure 4.**
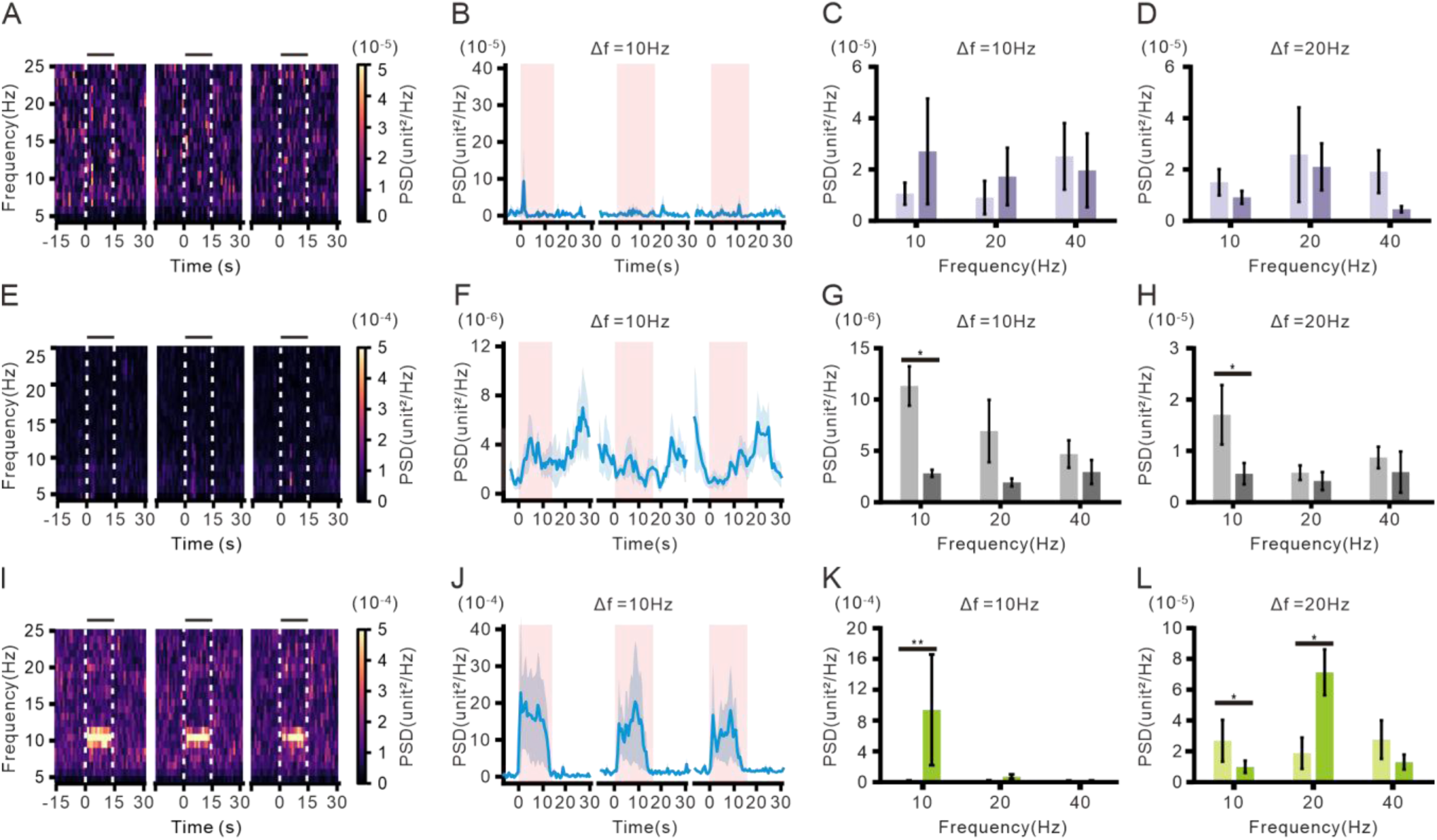
TIBS entrains envelope-frequency oscillations in a cell-type-specific manner. (**A**) Average time– frequency spectrogram of hippocampal CA2 neurons expressing AAV-hSyn-soma-jGCaMP8m during 10 Hz TIBS. Gray shaded regions indicate the stimulation periods. (**B**) Time course of the average 10 Hz PSD for the soma-targeted pan-neuronal recording; solid line, mean; light shaded area, SEM; pink shaded regions indicate the stimulation periods. (**C**) PSD at 10 and 20 Hz during the 5 s pre-stimulation baseline (light purple) and steady-state stimulation period (dark purple) under 10 Hz envelope TIBS (n = 6). (**D**) Similar to (C), but under 20 Hz envelope TIBS (n = 6). (**E**) Similar to (A), but with AAV-CaMKIIα-jGCaMP8m-WPRE for CaMKIIα-positive pyramidal neurons. (**F**) Similar to (B), but for the CaMKIIα-positive pyramidal recording. (**G**) Similar to (C), but for CaMKIIα-positive pyramidal neurons; baseline (light gray) and stimulation (dark gray) under 10 Hz envelope TIBS (n = 6). (**H**) Similar to (D), but for CaMKIIα-positive pyramidal neurons under 20 Hz envelope TIBS (n = 6). (**I**) Similar to (A), but with AAV-mDlx-jGCaMP8m-WPRE for GABAergic interneurons. (**J**) Similar to (B), but for mDlx-positive interneurons. (**K**) Similar to (C), but for mDlx-positive interneurons; baseline (light green) and stimulation (dark green) under 10 Hz envelope TIBS (n = 8). (**L**) Similar to (D), but for mDlx-positive interneurons under 20 Hz envelope TIBS (n = 6). *p < 0.05, **p < 0.01, ***p < 0.001, Wilcoxon signed-rank test.

At 10 Hz envelope stimulation, the soma-restricted pan-neuronal (hSyn-soma) recording showed a moderate elevation of PSD at 10 Hz of approximately 2-fold above baseline during stimulation (Figure 4A-C). The CaMKII-positive pyramidal population showed no envelope-frequency entrainment, and the PSD across all bands instead decreased during stimulation (Figure 4E-G). By contrast, the strongest envelope-frequency entrainment was observed in the mDlx-positive GABAergic interneuron population, where the PSD at 10 Hz increased approximately 60-fold above baseline (Figure 4I-K).

At 20 Hz envelope stimulation, the same qualitative pattern was observed. The hSyn-soma recording showed a modest elevation of PSD at 20 Hz above baseline (Figure 4D), the CaMKII-positive pyramidal population showed no envelope-frequency entrainment (Figure 4H), and the mDlx-positive interneurons again showed selective entrainment at the envelope frequency, with PSD at 20 Hz approximately tripling above baseline (Figure 4L). In the same mDlx recordings, the PSD at 10 Hz simultaneously decreased to about 37 % of baseline.

### 2.6 Local CA2 suppression does not extend to pyramidal axons or upstream cortical inputs

We next tested whether the pyramidal suppression extends to upstream cortical projections using retrograde AAV labelling. The AAVrg-CamKIIa-jGCaMP8m-WPRE was unilaterally injected into the CA2, driving GCaMP8m expression in upstream entorhinal/ectorhinal (Ent/Ect) pyramidal neurons that project to CA2 and additionally infecting a population of CA2 pyramidal neurons at the injection site (Figure 5A-B). With the optic fiber positioned in CA2, the photometry signal reflected the locally infected pyramidal neurons and the axon terminals of upstream cortical projections. This mixed signal showed only a mild monotonic decline from 1.94 ± 0.57 at 10 Hz to 1.09 ± 0.36 at 70 Hz, remaining positive across all envelope frequencies (Figure 5C-D), in marked contrast to the suppression below baseline observed at the same site with pure CaMKIIa- GCaMP8m (compare Figure 2H).

**Figure 5.**
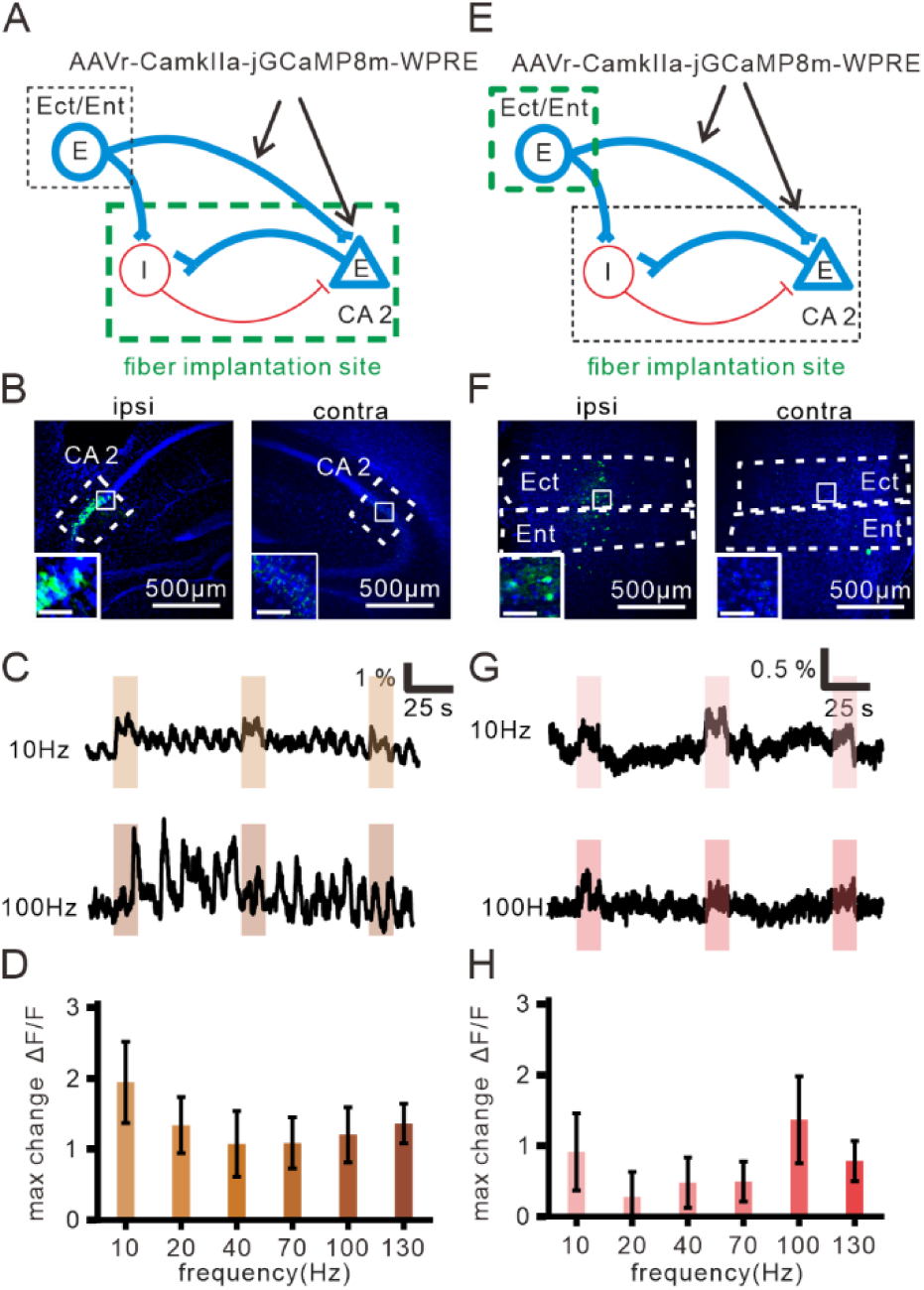
Retrogradely labelled CA2-projecting neurons show no frequency-dependent suppression. AAVrg-CaMKIIα-jGCaMP8m-WPRE was unilaterally injected into the right CA2 in all experiments. (**A**) Schematic of retrograde labelling and fiber photometry recording in hippocampal CA2. Red and blue cells denote inhibitory and excitatory neurons, respectively; thick lines denote sites of viral expression; dashed boxes denote brain regions; thick green boxes denote the recording location. (**B**) Representative fluorescence images of viral expression in the contralateral (left) and ipsilateral (right) hippocampal CA2. Insets show magnified views of the white-boxed regions. Green, GCaMP8m; blue, DAPI. Scale bars: main, 500 μm; insets, 80 μm. (**C**) Representative calcium traces recorded in CA2 during 10 Hz (upper) and 100 Hz (lower) envelope TIBS. Brown shaded regions indicate the stimulation periods. (**D**) Quantification of the absolute peak calcium response at different envelope frequencies (n = 9). (**E**) Similar to (A), but with fiber photometry recording in the ipsilateral Ect/Ent cortex. (**F**) Similar to (B), but with representative fluorescence images from the ipsilateral Ect/Ent. (**G**) Similar to (C), but recorded from the Ect/Ent; red shaded regions indicate the stimulation periods. (**H**) Similar to (D), but for the Ect/Ent recordings (n = 8). For all quantified data, the response was defined as the absolute peak ΔF/F, corresponding to the larger magnitude of either the maximum positive or minimum negative deflection during stimulation. Bar shade denotes envelope frequency, with darker shades indicating higher frequencies. *p < 0.05, **p < 0.01, ***p < 0.001, Repeated-measures ANOVA followed by Holm’s post-hoc test.

To isolate the upstream cell-body signal, we repeated the AAVrg-CamKIIa-jGCaMP8m-WPRE injection into CA2 in a separate cohort and implanted the optic fiber in the ipsilateral ectorhinal cortex (Figure 5E-F; AP −1.94 mm, ML +3.90 mm, DV −3.40 mm). Photometry at this upstream cortical site showed a small positive response to TIBS that was essentially flat across envelope frequencies (ΔF/F 0.28–1.36 % across 10–130 Hz; Figure 5G-H). Together, these data indicate that the strong frequency-dependent suppression of CA2 pyramidal somata observed locally does not produce a corresponding modulation of axon-terminal or upstream cell-body activity.

## 3 Discussion

In this study, we used cell-type-specific fiber photometry combined with immunostaining to dissect how dorsal CA2 of the mouse hippocampus responds to TIBS across a wide range of envelope frequencies. We found that envelope frequency operates as a tunable parameter that biases hippocampal E/I balance: low envelope frequencies activate both glutamatergic pyramidal and GABAergic interneuron populations, whereas envelope frequencies above approximately 20 Hz suppress pyramidal somatic activity below baseline while continuing to recruit GABAergic interneurons, with the divergence between the two populations reaching a maximum at 100 Hz. Retrograde labelling of upstream cortical projections to CA2 indicated that this E/I crossover is generated locally within the dorsal hippocampal circuitry, and FFT analysis of pan-neuronal and cell-type-specific photometry showed that only mDlx^+^ GABAergic interneurons exhibited robust envelope-frequency entrainment, accompanied by a concurrent suppression of endogenous lower-frequency rhythmic activity.

Among the four hypotheses for high-frequency DBS-induced suppression (Chiken & Nambu, 2016), our data support hypothesis (i), selective recruitment of local GABAergic interneurons that override pyramidal output, because mDlx-positive interneuron activity rises monotonically to peak at 100 Hz envelope frequency while CaMKII-positive pyramidal activity falls below baseline from 20 Hz onward, with maximum suppression at the same 100 Hz envelope frequency. In contrast, our data are inconsistent with the other three hypotheses. Hypothesis (ii), presynaptic synaptic depression, would predict elevated upstream activity as vesicles deplete, but retrogradely labelled upstream cortical projections showed frequency-invariant activity. Hypothesis (iii), direct depolarisation block, predicts uniform attenuation of all local populations including the interneurons themselves and is ruled out by the observed simultaneous interneuron activation at high frequency. Hypothesis (iv), information-lesion-type disruption, requires that target output remain present with altered temporal patterning, which is incompatible with the active reduction of pyramidal somatic Ca²⁺ signal below baseline observed here. This cell-type selectivity is biophysically consistent with the differential ion-channel composition of the two populations: PV-positive interneurons express Nav1.1 sodium channels together with fast Kv3-family potassium channels that support sustained gamma-frequency firing (Rudy & McBain, 2001; Ogiwara et al., 2007), whereas pyramidal neurons express Nav1.6 with slower delayed-rectifier kinetics (Kole et al., 2008). This intrinsic difference provides the first in vivo cell-type-resolved test of the biophysical prediction that envelope demodulation depends on active ion-channel rectification and should therefore differ across cell types (Mirzakhalili et al., 2020).

The cell-type and frequency-tuning patterns reported here have direct parallels in clinical deep brain stimulation of hippocampal-related targets. 130 Hz envelope TIBS applied to the mouse hippocampus suppresses pathological fast ripples and interictal epileptiform discharges in a kindling model of temporal lobe epilepsy (Acerbo et al., 2022); our data provide a cellular mechanism, since at 100 Hz and above pyramidal somatic activity in CA2 is strongly suppressed while PV⁺ interneurons remain robustly engaged, providing the sustained inhibitory drive that would attenuate pathological pyramidal output. Similar high-frequency choices are used in clinical DBS of hippocampal-circuit targets, including 145 Hz anterior thalamic DBS for drug-resistant focal epilepsy (Fisher et al., 2010) and 130 Hz fornix DBS investigated for memory impairment in Alzheimer’s disease (Lozano et al., 2016). At the opposite end of the envelope-frequency spectrum, 5 Hz envelope TIBS of the human hippocampus modulates BOLD signals and improves associative memory in healthy volunteers (Violante et al., 2023), corresponding to the low-Δf regime in which our data show broad activation of both excitatory and inhibitory populations. Together, these comparisons support a frequency-as- knob framework in which the envelope frequency of TIBS can be tuned to bias hippocampal output toward net activation (low Δf) or net inhibition (high Δf), paralleling the frequency-dependent effects of conventional DBS at hippocampal-related targets.

Several limitations of the present study should be acknowledged. First, all photometry recordings were performed under mild isoflurane anaesthesia to reduce spontaneous-activity interference and movement artefacts and to stabilise the optical interface; awake recordings will be needed to confirm that the frequency-dependent E/I shift generalises to behaviourally relevant states. Second, the kinetics of jGCaMP8m limit the reliable detection of envelope-locked oscillations to envelope frequencies of 20 Hz or below; the cell-type differences in mean ΔF/F observed up to 130 Hz indicate that the E/I shift is preserved at higher envelope frequencies, but direct spectral evidence of envelope-locked oscillations at gamma-band frequencies will require complementary electrophysiological recordings. Third, although the c-fos and parvalbumin co-staining corroborate PV^+^ interneurons as the principal active inhibitory subpopulation at high envelope frequencies, the mDlx promoter labels GABAergic interneurons across all subtypes. Fourth, the present framework is expected to extend to other hippocampal-like targets that share glutamatergic-pyramidal and PV- positive-interneuron architecture, but its applicability to regions with different cytoarchitecture, such as the striatum, will require direct testing. Finally, the present study documents how TIBS modulates neuronal activity but does not test behavioural consequences; future studies combining cell-type-specific photometry with hippocampal-dependent behaviour will be required to link these cellular findings to TIBS-induced cognitive and clinical effects.

## 4 Methods

### 4.1 Animals

All animal procedures were approved by the Institutional Animal Care and Use Committee of National Yang Ming Chiao Tung University (IACUC NO.112016) and were conducted in accordance with the Guide for the Care and Use of Laboratory Animals of NYCU. Adult (8 to 20 weeks old) male C57BL/6 mice and male Thy1- GCaMP6s transgenic mice (C57BL/6J-Tg(Thy1-GCaMP6s)GP4.3Dkim/J; Dana et al., 2014) were used. A total of 66 mice were included across all experiments; per-cohort sample sizes are reported in the corresponding figure legends. C57BL/6 mice were purchased from BioLASCO Taiwan Co. Ltd. and Thy1- GCaMP6s mice from Jackson Laboratory (stock no. 024275). Animals were group-housed in individually ventilated cages on a 12 h light/dark cycle at the NYCU Laboratory Animal Center, with food and water available ad libitum.

### 4.2 Viral constructs

Recombinant adeno-associated viruses (AAVs) were used to drive cell-type-specific expression of jGCaMP8m in hippocampal CA2 or in upstream entorhinal / ectorhinal cortex neurons projecting to CA2. Five viral constructs were used: hSyn (pan-neuronal), hSyn-soma (soma-restricted pan-neuronal), CaMKIIα (excitatory pyramidal), mDlx (GABAergic interneurons), and a retrograde AAVrg-CaMKIIα for upstream cortical projections. Viral titers were ≥1 × 10¹³ GC mL⁻¹. Full construct names and sources: pGP-AAV-syn- jGCaMP8m-WPRE (pan-neuronal; gift from the GENIE Project, Addgene plasmid #162375, viral prep #162375-AAV9; Zhang et al., 2020); AAV-hSyn-soma-jGCaMP8m (soma-restricted pan-neuronal; gift from Marianne Fyhn, Addgene plasmid #169257, viral prep #169257-AAV9; Grødem et al., 2023); AAV-CamKIIa- jGCaMP8m-WPRE (excitatory pyramidal; gift from Loren Looger, Addgene plasmid #176751, viral prep #176751-AAV9); AAV-mDlx-jGCaMP8m-WPRE (GABAergic interneurons; gift from Loren Looger, Addgene plasmid #176754, viral prep #176754-AAV9); AAVrg-CamKIIa-jGCaMP8m-WPRE (retrograde; from Addgene plasmid #176751, viral prep #176751-AAVrg).

### 4.3 Stereotaxic surgery and implantation

Mice were anesthetised with 1.5–2% isoflurane in oxygen and secured on a stereotaxic frame (71000, RWD) with body temperature maintained at 37.5 °C. AAV vectors were injected into the right CA2 (AP −1.94, ML +2.36, DV −2.0 mm from bregma; 0.5 μL at 0.1 μL min⁻¹). After one month of viral expression, four stainless-steel stimulating electrodes and one grounding electrode were implanted in the skull, together with an optic fiber (Φ200 μm core, 0.50 NA) targeting either dorsal CA2 (AP −1.94, ML +3.90, DV −1.8 mm) or the right ectorhinal cortex (AP −1.94, ML +3.90, DV −3.40 mm). For c-fos experiments without fiber photometry, only the stimulating and grounding electrodes were implanted at the same coordinates.

Ophthalmic ointment was applied to prevent corneal drying. Craniotomies were performed with a dental drill above the target coordinates. Each viral injection delivered 0.5 μL at 0.1 μL min⁻¹ using a microinjection syringe (7653-01, Hamilton) fitted with a 33-gauge needle (7803-05, Hamilton) and driven by a micropump (LEGATO130, Kd Scientific). The four stimulating electrodes were fixed on the skull at the following coordinates (AP, MLin mm, relative to bregma): (+2.20, +2.04), (+1.10, +2.04), (−4.40, +2.04), and (−5.50, +2.04). A grounding electrode was placed at (−1.94, −3.90). The fiber ferrule and electrodes were secured to the skull with dental resin (Super Bond, Sun Medical). For c-fos experiments without fiber photometry, the stimulating and grounding electrodes were secured with dental adhesive (3M Single Bond Universal Adhesive). Following surgery, Carprofen (5 mg kg⁻¹) was administered subcutaneously in the neck for post- operative analgesia. Mice were monitored daily until recovery. Photometry recordings began on the same day as fiber implantation when mice exhibited stable post-surgical condition; otherwise, recordings were postponed until the animals had fully recovered.

### 4.4 TIBS system and stimulation protocol

Temporal interference stimulation was delivered through a custom-built multi-channel current stimulator controlled by a LabVIEW-based interface. Two pairs of alternating currents with carrier frequencies f₁ = 2.00 kHz and f₂ = 2.00 kHz + Δf produced a low-frequency amplitude envelope Δf at the field intersection. Δf was set to 10, 20, 40, 70, 100, or 130 Hz in separate trials, with peak-to-peak current amplitude ≤2 mA per electrode pair, within safety thresholds for transcranial electrical stimulation (Cassara et al., 2025). Each stimulation block comprised a 2 s ramp-up, 10 s plateau, and 2 s ramp-down; blocks were separated by 1 min inter-stimulus intervals with envelope-frequency order randomised across animals and sessions. Three epochs per condition were averaged per animal. Control conditions comprised two negative controls activating one pair at a time (Pair 1 alone at 2.00 kHz; Pair 2 alone at 2.01 kHz) and a positive control delivering tACS at the envelope frequency between two distal electrodes from different pairs.

The custom-built stimulator comprised a signal input stage, a voltage-to-current (V-to-I) conversion circuit, four independent output channels, and a microcontroller unit (MCU). External stimulation waveforms were supplied by an Analog Discovery 3 (AD3, Digilent) or an equivalent signal generator and converted into stable stimulation currents through the V-to-I circuit. Stimulation parameters, including waveform type (sinusoidal or square), carrier frequency, current amplitude, and ramp settings, were controlled through a LabVIEW-based graphical user interface. The output current amplitude was adjustable in the range of approximately 0.1 to 3 mA per electrode pair, and the entire stimulator was powered by four 9 V batteries to provide a portable and electrically isolated stimulation platform.

### 4.5 COMSOL electric field simulation

To predict the electric-field distribution generated by TIBS in the mouse brain, we adapted a previously published finite-element mouse head model implemented in COMSOL Multiphysics (v6.3, AC/DC module), computing the modulation depth of the 10 Hz amplitude-modulated field in the z-direction (Ez). The simulated modulation-depth map was validated against c-fos immunostaining patterns following in vivo TIBS by thresholded overlap.

We adapted a three-dimensional finite-element mouse head model previously developed by our group (Chou et al., 2025), retaining the original tissue compartments, electrical conductivities, and electrode specifications. Several geometric refinements were introduced to improve anatomical correspondence with the Allen Mouse Brain Atlas: the head was modelled as an ellipsoid with semi-axes of 5, 6.5, and 3 mm, and the brain as an inner ellipsoid with semi-axes of 3.5, 5.5, and 2.2 mm. The coordinate system was redefined so that bregma corresponded to (0, 0, 0), placing the geometric centre of the brain ellipsoid at (AP, ML, DV) = (−2.6045, 0, −2.7487) mm. To correct residual geometric mismatch between the simplified ellipsoid model and the asymmetric morphology of the actual mouse brain, an empirical anatomical calibration was applied: the entire model was uniformly translated 0.4 mm laterally (right), 0.2 mm posteriorly, and 0.27 mm dorsally, so that the computational coordinate space aligned with the stereotaxic space used for experimental electrode placement. Two pairs of stimulating electrodes were modelled at the following coordinates (AP, ML, DV in mm, relative to bregma): (2.2, 2.04, −1.3), (1.1, 2.04, −0.7), (−4.4, 2.04, −0.2), and (−5.5, 2.04, −0.4). The ground electrode was placed at (−1.94, −3.9, −1). The two pairs were driven as 1.6 mA current sources at 2000 Hz and 2010 Hz, respectively. The modulation depth of the 10 Hz amplitude-modulated field in the z-direction (Ez) was defined as the difference between the upper and lower envelopes of the superposed time-domain signal at each mesh node. To validate the spatial specificity of the simulated field against biological activation, the simulated modulation-depth map was binarised across a series of candidate activation thresholds; for each threshold, the predicted activation region was compared with the c-fos-positive region, and the threshold yielding the highest spatial agreement with the biological measurement was selected as the optimal activation criterion.

### 4.6 Fiber photometry recording

Calcium activity was recorded using a dual-wavelength fiber photometry system (R810, Nanjing Zhike Biotechnology; Cheng et al., 2025) with alternating 470 nm (GCaMP excitation) and 410 nm (isosbestic control) illumination. Signals were acquired at 60 or 300 frames per second through a low-autofluorescence patch cord (Φ200 μm core, 0.50 NA). Recordings were performed under mild isoflurane anaesthesia (1.5% in oxygen) to minimise movement artefacts, with baseline fluorescence stability confirmed for 20 s before each recording. LED powers were set to 38.11% at 470 nm and 17.18% at 410 nm to balance signal-to-noise ratio against photobleaching. The TIBS stimulator was synchronised with the photometry system through a digital output signal, ensuring precise temporal alignment between stimulation and fluorescence acquisition.

Photometry was performed on the same day as fiber implantation for experiments requiring sequential recordings at multiple cortical depths within a single session, such as the Thy1-GCaMP6s spatial-profile recordings, in which the fiber was advanced through successive depths under the same anaesthesia. For all other experiments, recordings began after a post-operative recovery period of one day.

### 4.7 Calcium signal analysis

Raw fluorescence signals were processed offline in Python. Motion and bleaching correction was performed by least-squares fitting the 410 nm isosbestic channel to the 470 nm GCaMP channel to yield ΔF, and ΔF/F₀ was computed by normalisation to baseline F₀. Maximum, mean, and minimum ΔF/F values were extracted from each stimulation trial; the baseline epoch was defined as the interval from −10 to 0 s and the stimulation epoch as 2 to 12 s relative to stimulation onset, allowing for the indicator’s onset kinetics. Three epochs per condition were averaged per animal for subsequent statistical analyses. Stimulus-aligned traces were anchored to the digital event marker (“Input1*2*0;”) recorded with the TIBS stimulator and designated as t = 0 s. Processed measurements were exported as structured CSV files.

### 4.8 Fast Fourier transform and spectrogram analysis

Spectral analysis of ΔF/F traces was performed offline in Python. Time–frequency spectrograms were generated using a sliding-window fast Fourier transform (FFT) approach. A 5 s analysis window with a 0.5 s step size was applied across the recording, and the power spectral density (PSD) for each window was calculated using the same FFT-based PSD estimation as that used for the global spectral analysis. Spectral power was visualised in the 4–25 Hz frequency range to examine stimulation-related modulation of neural activity. PSD was expressed in a.u. Hz⁻¹ and quantified across three temporal epochs: (i) pre-stimulation baseline (5 s preceding stimulation onset), (ii) the stimulation period, and (iii) the full recording session. For frequency-specific analyses, the power at the stimulation envelope frequency was extracted from the spectrogram and tracked over time to characterise the temporal dynamics of TIBS-induced neural entrainment.

### 4.9 Histology and immunostaining

For c-fos experiments, mice were sacrificed 90 min after TIBS and perfused with 1× PBS followed by 4% paraformaldehyde (PFA). Brains were post-fixed in 4% PFA overnight at 4 °C and sectioned coronally at 60 μm using a vibratome. Sections were permeabilised, blocked, and incubated overnight with primary antibodies against c-Fos and parvalbumin (PV), followed by fluorescently conjugated secondary antibodies (2 h at room temperature) and mounted with DAPI-containing medium.

Sections were washed three times with PBS for 5 min each and permeabilised in 2% (v/v) Triton X-100 (Sigma-Aldrich) for 20 min. Endogenous peroxidase activity and tissue background were quenched with a 10 min incubation in a mixture of 2% Triton X-100, 30% H₂O₂, and methanol. After three additional PBS washes, sections were blocked in 3% normal goat serum in PBS for 60 min at room temperature (20 to 26 °C). Primary antibodies were diluted in PBS containing 1% normal goat serum and 2% Triton X-100 and applied overnight at 4 °C: rabbit anti-c-fos (9F6 #2250, Cell Signaling Technology, 1:500) and mouse anti-parvalbumin (P3088, Sigma-Aldrich, 1:1000). After three PBS washes, sections were incubated for 2 h at room temperature with the appropriate secondary antibodies (all at 1:500): goat anti-rabbit Alexa Fluor 594 (ab150080, Abcam), goat anti-rabbit Alexa Fluor 488 (ab150077, Abcam), goat anti-mouse Alexa Fluor 488 (ab150113, Abcam), and goat anti-mouse Alexa Fluor 594 (ab150116, Abcam). Sections were mounted on glass slides with Fluoroshield containing DAPI (GTX30920, GeneTex).

### 4.10 Image acquisition and quantification

Fluorescence images were acquired on an upright fluorescence microscope (SS-1000-00, Scientifica) equipped with an LED source, sCMOS camera, and DAPI/GFP/TexasRed filter cubes. Quantification of c-Fos and PV immunoreactivity was performed in ImageJ / Fiji using a custom batch-processing macro that segmented c-fos⁺ signals with local thresholding and ellipse fitting, then required overlap with DAPI-defined nuclei; the same pipeline identified PV⁺ and c-fos⁺/PV⁺ double-labelled cells. Counts were obtained by an experimenter blind to the stimulation condition. Fiber and electrode placement was verified in every animal; mice with off-target placement were excluded from analysis.

Fluorescence images were acquired on an upright fluorescence microscope (SS-1000-00, Scientifica) equipped with an LED light source (pE300, CoolLED), a Hamamatsu C13440 camera, and filter cubes (39000, 19008, and 31002, Chroma Technology) for DAPI, GFP, and TexasRed channels. In ImageJ / Fiji, background was attenuated using a rolling-ball algorithm (radius = 50 pixels) followed by Gaussian blur (σ = 2 pixels). c-Fos-positive signals were segmented using the Phansalkar local-thresholding method and converted to normalised elliptical regions of interest (ROIs) by ellipse fitting, ensuring that morphological variation did not bias the count. Cells were defined as c-fos⁺ only when their ROI spatially overlapped with a DAPI-defined nucleus, as determined by binary AND mask intersection. The same pipeline was applied to identify PV⁺ and c-fos⁺/PV⁺ double-labelled cells. Counts were obtained from 1 non-adjacent section per mouse and 6 mice per condition and were extracted within predefined anatomical ROIs and exported as structured CSV files.

### 4.11 Statistical analysis

Statistical analyses were performed in JASP (v0.18.3.0, JASP Team). Data are reported as mean ± SEM (Standard Error of Mean). Repeated-measures ANOVA with Holm’s post-hoc test was used for within-cell-type comparisons across envelope frequencies; the Wilcoxon signed-rank test was used for paired comparisons where non-parametric testing was required. The sample size n refers to individual animals unless otherwise specified. All tests were two-tailed; statistical significance was set at p < 0.05, and exact p values are reported in the figures and figure legends.

## Author Contributions

**Yi-Cheng Fang:** Data Curation, Methodology, Investigation, Formal analysis, Visualization, Validation, Writing –review & editing. **Lin Chou:** Methodology, Software (TIBS device). **Yao-Yi Tseng:** Software (COMSOL simulation), Formal analysis. **Po-Hsun Chu:** Methodology (TIBS device). **Yi-Man Fang:** Methodology (TIBS device). **Hai-Yin Chen:** Methodology (TIBS device). **Yu-Te Liao:** Supervision, Methodology, Resources, Funding acquisition. **Chih-Hsien Huang:** Supervision, Methodology, Formal analysis. **Po-Han Chiang:** Supervision, Conceptualization, Methodology, Software, Formal analysis, Investigation, Visualization, Funding acquisition, Project administration, Writing – Original draft, Writing – Review & Editing.

## Acknowledgements

We thank Prof. Tsai-Wen Chen (National Yang Ming Chiao Tung University) for kindly providing the Thy1- GCaMP6s transgenic mice. We are also grateful to Prof. Albert C. Yang (National Yang Ming Chiao Tung University) for insightful discussion of the study and for assistance with funding acquisition.

## Data Availability Statement

The data that support the findings of this study are available from the corresponding author upon reasonable request.

## Ethics Statement

All animal procedures were approved by the Institutional Animal Care and Use Committee (IACUC) of National Yang Ming Chiao Tung University under protocol number IACUC NO.112016 (PI: Prof. Po-Han Chiang), and were conducted in accordance with the Guide for the Care and Use of Laboratory Animals of NYCU. The experimental design, procedures, and reporting adhered to the ARRIVE 2.0 guidelines (Percie du Sert et al., 2020). Consistent with the 3Rs principles of Replacement, Reduction, and Refinement, sample sizes were minimised while retaining sufficient statistical power, and postoperative analgesia and daily monitoring were provided to reduce animal suffering. Full details of housing conditions, surgical procedures, and euthanasia are provided in the Methods.

## Declaration of generative AI and AI-assisted technologies in the manuscript preparation process

During the preparation of this work, the author used Claude (Anthropic), Gemini, and ChatGPT for assistance with manuscript editing and code development for data analysis. The authors reviewed and verified all outputs and take full responsibility for the content of this work.

